# Consistent behavioral syndrome across seasons in an invasive freshwater fish

**DOI:** 10.1101/2020.03.03.974998

**Authors:** Juliane Lukas, Gregor Kalinkat, Friedrich Wilhelm Miesen, Tim Landgraf, Jens Krause, David Bierbach

## Abstract

Understanding the linkage between behavioral types and dispersal tendency has become a pressing issue in light of global change and biological invasions. Here, we explore whether dispersing individuals exhibit behavioral types that differ from those remaining in the source population. We investigated a feral population of guppies (*Poecilia reticulata*) that undergoes a yearly range shift cycle. Guppies are among the most widespread invasive species in the world, but in temperate regions these tropical fish can only survive in winter-warm freshwaters. Established in a thermally-altered stream in Germany, guppies are confined to a warm-water influx in winter, but can spread to peripheral parts as these become thermally accessible. We sampled fish from the source population and a winter-abandoned site in March, June and August. Fish were tested for boldness, sociability and activity involving open-field tests including interactions with a robotic social partner. Guppies differed consistently among each other in all three traits. Behavioral trait expression in the source population differed across seasons, however, we could not detect differences between source and downstream populations. Instead, all sampled populations exhibited a remarkably stable behavioral syndrome between boldness and activity despite strong seasonal changes in water temperature and associated environmental factors. We conclude that random drift (opposed to personality-biased dispersal) is a more likely dispersal mode for guppies, at least in the investigated stream. In the face of fluctuating environments, guppies seem to be extremely effective in keeping behavioral expressions constant, which could help explain their successful invasion and adaptation to disturbed habitats.

## Introduction

Consistent behavioral differences among individuals (e.g. behavioural types or animal personality; Réale et al. 2007) seem to be a ubiquitous biological feature (Bell et al. 2009; Bierbach et al. 2017) that was proposed to have substantial ecological and evolutionary importance (Sih et al. 2012; Wolf and Weissing 2012). For one, behavioural types are thought to be relevant in determining success (Chapple et al. 2012; Carere and Gherardi 2013; Canestrelli et al. 2015) and impact of biological invasions (Cote, Clobert, et al. 2010; Fogarty et al. 2011; Juette et al. 2014), which is becoming an ever more pressing issue in the light of global change and ongoing translocations of species among habitats (Wong and Candolin 2015; Seebens et al. 2017; Seebens et al. 2018).

It is assumed that individuals that invade novel habitats are a non-random subset of their source populations (Shine et al. 2011; Sih et al. 2012; Juette et al. 2014; Canestrelli et al. 2015; Spiegel et al. 2017). These invaders may share certain phenotypic and life-history characteristics that are associated with their behavioral types (Juette et al. 2014). Behavioral types of invading individuals may determine whether they are transported (by humans) or move (by themselves) into new environments (first invasion stage; Lockwood et al. 2011), whether they can survive and establish a non-native population (second stage), and whether they may spread to new habitats thereafter (third stage). Behavioral types shown to be relevant in this regard are boldness (Cote et al. 2011; Myles-Gonzalez et al. 2015), aggression (Duckworth and Badyaev 2007; Hudina et al. 2015), activity (Brown and Irving 2014; Thorlacius et al. 2015) as well as sociability (Cote, Fogarty, et al. 2010; Cote et al. 2011; Rasmussen and Belk 2012). Further, it was predicted that individuals carrying correlated suites (i.e., behavioral syndromes, Sih et al. 2012) of bold, aggressive, active and asocial types may predominantly be found at the invasion front (Duckworth and Badyaev 2007; Cote, Fogarty, et al. 2010; Cote et al. 2011; Juette et al. 2014). However, such spatially and/or temporally assorted compositions of behavioral types among populations have only rarely been investigated along naturally occurring invasion gradients.

The difficulty to study invasion phenomena in the wild is their random and unpredictable occurrence (Lockwood et al. 2011), which makes it nearly impossible to follow active invasion fronts. In the rare cases where range expansions could be tracked in real time (e.g., western bluebirds: Duckworth and Badyaev 2007; African jewelfish: Lopez et al. 2012; signal crayfish: Hudina et al. 2015; round goby: Thorlacius et al. 2015; cane toad: Gruber et al. 2017), the invasion origin is often unknown or unclear due to multiple introduction events. Thus, we need to identify study systems that allow us to investigate dispersal across sensible temporal and spatial scales. Thermally altered freshwater systems (TAFs) in the temperate zones provide fruitful conditions to study this phenomenon. Here, natural and/or anthropogenic warm water influxes create unique temperature refuges that allow non-native species (especially those of tropical origin) to establish populations (Langford 1990; Jourdan et al. 2014). While ambient water temperatures drop in winter (e.g., less than +1°C in the river Rhine; Jourdan et al. 2014), areas close to constant warm water influx will remain suitable for non-native tropical species to survive. Once peripheral areas become thermally suitable again in warmer seasons, certain individuals from the sources may disperse into these areas. Such a scenario enables us to investigate the role of behavioral types during *in situ* dispersal of non-native species from a single, well-determined source into new (i.e. winter-abandoned) habitats.

Under a personality-biased dispersal hypothesis, where individuals with a certain set of personality traits are assumed to leave the source population to colonize new habitats, we predicted that *(i)* the source population’s average personality traits should change over the course of the year due to the drain of the emigrating individuals, *(ii)* newly founded populations should differ in their average personality traits compared to their source populations, and *(iii)* behavioral variation should decrease both in the source as well as in the newly founded population due to the non-random migration patterns. In addition, *(iv)* population-wide correlations among personality traits (behavioral syndromes) may emerge or disappear because of assumed non-random migration patterns.

In Germany, tropical fishes have established populations in several artificially heated stream systems (Lukas, Kalinkat, et al. 2017), the most investigated being the Gillbach near Cologne (see Fig. 1a; Jourdan et al. 2014; Lukas, Jourdan, et al. 2017; Kempkes et al. 2018). To estimate seasonal fluctuations in temperature-associated abiotic and biotic conditions within the sampled range of the stream, we continuously recorded water temperatures with submerged data-loggers. We sampled feral guppies (*Poecilia reticulata*) from the Gillbach during different seasons (temporal scale: spring, early summer, late summer) and, if possible, at different sites (spatial scale: core of warm water influx versus more peripheral downstream area; see Fig. 1a) and repeatedly assessed behavioral types in terms of boldness, sociability and activity in a laboratory setting.

**Figure 1:**
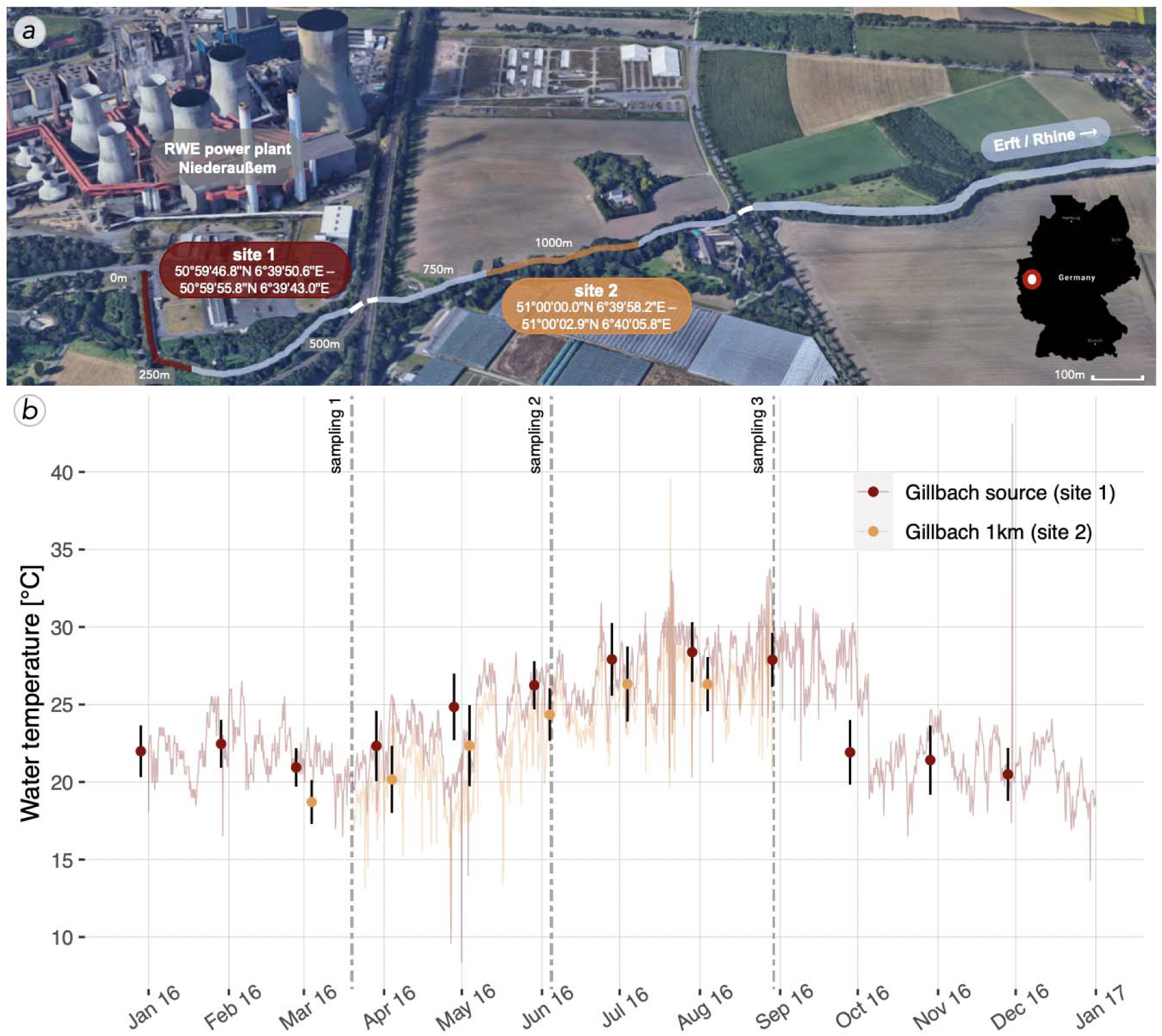
The Gillbach stream and its altered temperature regime. **a)** Warm water discharge from a power plant feeds into the Gillbach (map created using © 2019 Google Earth with data from 2009 GeoBasis-DE/BKG) and generates a thermal gradient above ambient water temperatures that is preserved for several kilometers (site 1 – site 2). **b)** 4-hourly water temperature* with error bars indicating monthly means ± SD for each site (site 1 – Gillbach source: red; site 2 – one kilometer downstream: orange). Dotted lines correspond to the three sampling occasions. * Note that due to a loss of loggers, continuous monitoring at site 2 was interrupted (no data from 01.01.2016 – 19.03.2016 nor from 29.08.2016 – 31.12.2016).

## Material& Methods

### Study site & Sampling

The Gillbach (Rhine catchment, Germany) has been under the influence of thermal pollution and anthropogenic modifications for over six decades. Its headwaters are exclusively fed by the warm water discharge of the brown coal power plant Niederaußem (Fig. 1a). This creates a temperature regime that is highly dependent on the volume and temperature of the plant’s water discharge. Generally exhibiting above ambient water temperatures (Fig. 1b), the Gillbach harbors several established populations of non-native fish, invertebrate and plant species, many of which are of tropical or subtropical origin (see Supplementary material Appendix 1, Table S1 for a list of native and non-native fish species found in the Gillbach).

Here, feral guppies (*Poecilia reticulata*) are highly abundant and as successful global invaders (Deacon et al. 2011) they are well suited to investigate the expression of behavioral traits across time and space. Sampling was conducted at two sites (∼300m transects, see Fig. 1a) in March, June and August 2016. The Gillbach’s guppy population density increased from March to June and decreased in late summer (Supplementary material Appendix 2, Table S3). This extreme population decline was likely predation-induced as densities of convict cichlids (*Amatitlania nigrofasciata*) and European chub (*Squalius cephalus*) also increased during this period (Supplementary material Appendix 1, Table S1). Sampling commenced until sufficient numbers for behavioral testing were reached (i.e., ∼40 adult individuals). Captured fish were transferred into well-aerated coolers and kept separately by sampling site. To reduce a possible sampling bias of certain behavioral types, we collected specimens by seine and dip netting and actively drove fish into the net as recommended by Biro & Dingemanse (2009). While we acknowledge some size selectivity against juvenile guppies introduced by the 2 mm mesh size, our population sample was representative of adult body size variation (see Fig. 3b), thus avoiding indirect bias of behavioral traits that are correlated with size (Polverino et al. 2016).

**Figure 2:**
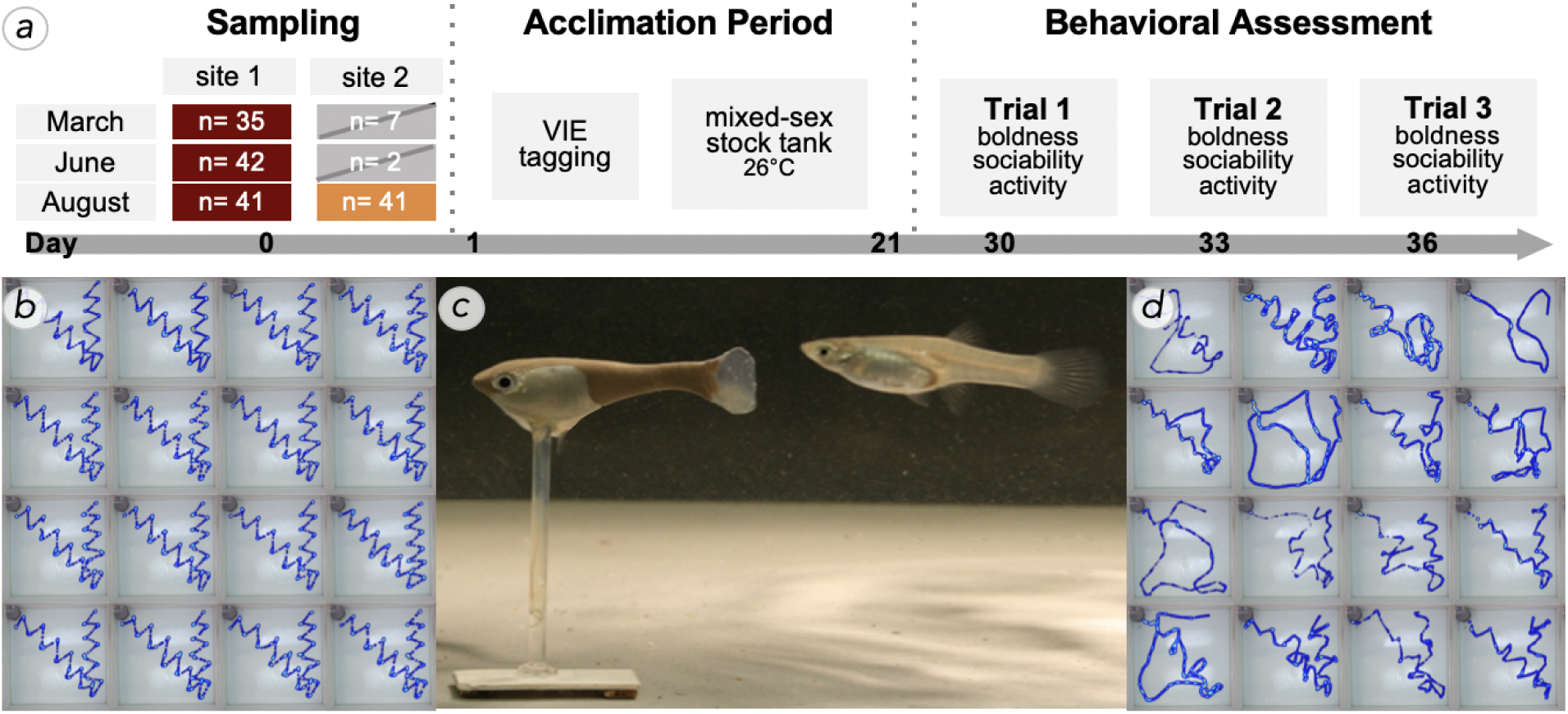
Behavioral testing with robotic fish. **a)** Test fish were collected three times and, when possible, at two sites (site 1 and 2, see Fig. 1a). After an initial 30-day acclimation period, fish were tested a total of three times (trial I – III, sample sizes indicate tested individuals, for total catch numbers see Supplementary material Appendix 2, Table S3). **b-d)** During trials, the fish replica was moved remotely through the test tank (c). Once the focal individual had emerged from its shelter, the biomimetic robot initiated its predefined path (b) and the fish’s swimming trajectory was recorded (d).

**Figure 3:**
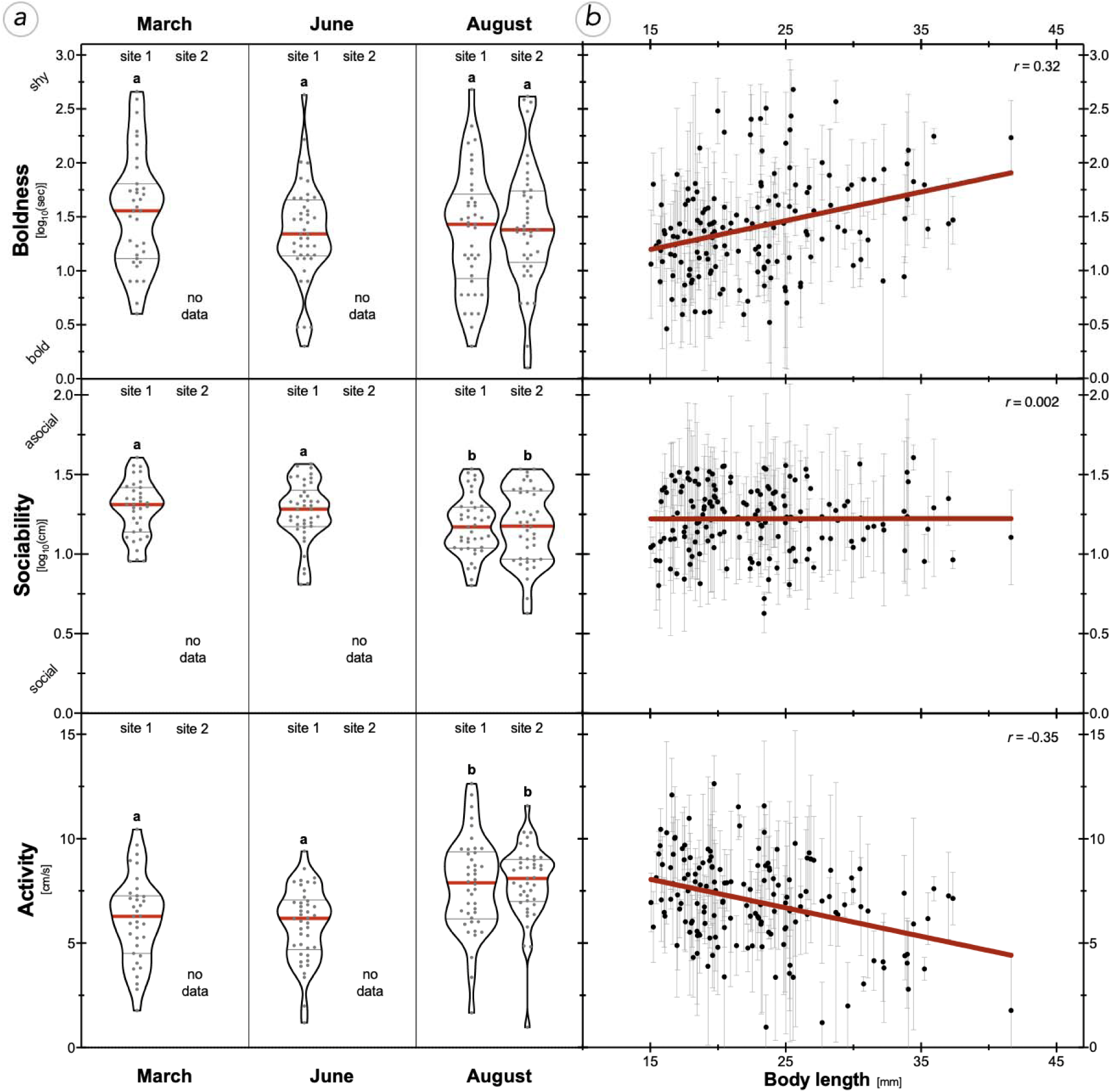
Average personality traits across temporal and spatial scale. ***(a)*** Median trait scores (and IQRs) for boldness* (i.e., time to emerge from shelter), activity (i.e., velocity during open field exploration) and sociability* (i.e., distance to biomimetic robot). Different letters depict significant differences based on post-hoc LSD tests after global linear mixed models (*p* < 0.05). ***(b)*** Mean traits scores (± SD) in relation to body size (standard length; *n*=159). To visualize the relationships, we show linear regressions lines (red) calculated from individual means (see main text for significance levels and Pearson’s *r*). * Note that low scores of boldness and sociability indicate high trait expression.

Water temperatures at the source (site 1) and one kilometer downstream (site 2) were monitored using HOBO data loggers (Onset Computer Corporation, Bourne, MA, USA) for a period of 12 months (Fig. 1b). For the period of sampling and behavioral testing (March – August 2016), the Gillbach experienced a thermal gradient of ∼2°C/km (mean ± SD, site 1: 25.53°C ± 3.21°C; site 2: 23.49°C ± 3.4°C; Fig. 1b). However, we expect the temperature difference between sites to be even more pronounced in colder months (i.e. decline of 3.5°C/km in November (Klotz et al. 2013); 4.7°C/km in February (Jourdan et al. 2014)). With temperatures below the population’s critical minimum temperature (∼12°C; Jourdan et al. 2014), survival of guppies outside the core area during winter is highly unlikely. Under similar conditions in June (∼1.5°C/km gradient), a previous study (Emde et al. 2016) found diet composition of the non-native predatory cichlid *Amatitlania nigrofasciata* to vastly differ between sites, with cichlids exerting more piscivorous predation pressure at peripheral sites.

### Behavioral assessment

Fish collected for behavioral testing were transferred to the laboratory at Humboldt University of Berlin, Germany. Fish were kept in one holding tank (270 liters), but were separated into mesh-divided compartments by collection time and site (‘populations’ hereafter). Each compartment (24 cm × 58 cm × 40 cm L×W×H) contained artificial java moss and natural gravel. Females and males were housed communally. Water was maintained at 26°C and kept under continuous aeration and filtration. Fish were held under diurnal lighting (12:12h light:dark cycle) and fed commercial flake food (TetraMin® Tetra GmbH) twice a day. Feeding stopped 12 hours before experiments and recommenced afterwards. To acclimate the fish to laboratory conditions and reduce potential short-term effects of the different sampling habitats on behavioral trait expression, no behavioral testing was conducted for at least 30 days (see Fig. 2a for detailed experimental design). One week prior to testing, adults of suitable size (SL>15 mm) were marked using visible implant elastomer tags (VIE; Northwest Marine Technology Ltd, Shaw Island, WA, USA) to allow for individual identification throughout the experiment. To be mindful of capture biases towards a specific behavioral type, we used all fish within a compartment for testing.

We tested a total of 159 guppies (95 females, 64 males; for population-specific sample sizes see Fig. 2a). Test fish were introduced to an open field test arena (88 × 88 cm, 75 mm water depth) filled with aged tap water kept at 26°C. To reduce disturbing stimuli, the arena was enclosed in black, opaque plastic and illuminated with artificial light reproducing the daylight spectrum. Behavioral observations were recorded via a camera (Ximea 4K USB 3.1 camera) mounted above the arena. Position information was extracted using EthoVision 10 XT software (Noldus Information Technology, Wageningen, Netherlands).

#### (1) Boldness

Latency to emerge from a shelter was used as a measure of boldness with a low latency time indicating high levels of boldness. Fish were introduced into a grey opaque shelter box (Ø 100 mm; top left of each image in Fig. 2b and d) and allowed to acclimate. After one minute, a sponge was removed from the entrance (40mm x 25mm), allowing the fish to emerge and explore the test arena. A fish was scored as having emerged when its full body was visible outside of the shelter. If fish did not emerge after five minutes, the lid of the shelter box was removed. Fish that failed to emerge within eight minutes were given the maximum score of 480 seconds and the box was removed entirely.

#### (2) Sociability

A three-dimensional fish replica was stationed 1 cm away from the shelter entrance. The replica, which resembled a large-sized guppy female (SL=30mm; left fish in Fig. 2c), could be moved by a robot via a magnetic base (see Landgraf et al. 2016; Bierbach et al. 2018 for details). Upon the fish’s emergence (or the removal of the shelter box), the biomimetic robot started to move through the arena in a standardized sequence (Fig. 2b) imitating a conspecific. We calculated the mean distance the fish kept to the biomimetic robot as a proxy of sociability with lower values indicating higher levels of sociability. Consequently, fish that followed the robot closer were assumed to be more sociable (see Bierbach et al. 2018).

#### (3) Activity

After completion of the robotic fish’s predefined path, the replica was removed from the arena by hand and once the fish resumed normal swimming behavior, it was allowed to explore the arena. As a measure of activity, we calculated the mean velocity with which the fish explored the open arena during 3 minutes.

To minimize handling and exposure to stress (Animal Behaviour 2020), all three traits were assessed within a single behavioral assay. After the first behavioral assay, we recorded the body length of each individual. Fish were transferred back to their respective compartment and allowed 72 hours resting before being retested following the same protocol. To reduce acclimatization effects, we only assessed short-term repeatability by testing fish a total of three times (randomized order) within the span of a week. All reported experiments comply with the current German law approved by LaGeSo (G0117/16 to Dr. D. Bierbach).

### Statistical analysis

To test for differences in average trait expression over a temporal and/or spatial scale, we used linear mixed models (LMM, *GENLINMIXED* procedure; SPSS v. 25.0, IBM Corp., Armonk, NY, USA) with each behavioral trait (boldness, sociability, activity) as a separate response variable. ‘Boldness’ and ‘sociability’ were log10-transformed to fulfil normality assumptions. All models included sample population as the fixed effect of interest (four levels: March – site 1, June – site 1, August – site 1, August – site 2). The low sample size collected at site 2 during March and June (*n*=7 and *n*=2, respectively; see Fig. 2a) made these estimates uncertain. After verifying that there were no substantial differences found when running the analyses with and without the above population samples, these estimates were excluded from final analyses. Behavior in poeciliids is often size- and/or sex-dependent (Brown and Braithwaite 2004; Harris et al. 2010), thus we included standard length (SL) as a covariate and ‘sex’ as an additional fixed factor. To account for the repeated testing of individuals, we further included ‘trial’ as a fixed factor and ‘fish ID’ as a random factor. Post-hoc LSD tests were employed in case fixed factors were found to have a significant effect. For the covariate ‘body size’, we also performed Pearson’s correlations to better evaluate the relationship.

(Co)Variance structure was nested within ‘population’, which enabled us to calculate the total behavioral variance as well as within- and among-individual variance components in each population. Based on this, we calculated behavioral repeatability for each of the four populations as a measure of consistency. We assumed that repeatability and/or variance estimates differed among populations when 95% confidence intervals (CIs) were not overlapping (see Nakagawa and Schielzeth 2010).

To evaluate the existence of behavioral syndromes (i.e. correlation among behavioral traits) within our populations, we used linear regression models to test for significant correlation between each trait pair. In order to minimize the effects of mean differences in behavioral traits on trait correlation, we mean-centered each behavioral trait within ‘trial’ and ‘population’ (z-transformation). Each model included one trait as the dependent variable and another as covariate. We further compared whether trait correlation differed among populations (i.e., difference in slopes termed ‘covariate nested within population’). Visualizations were done with GraphPad Prism (version 8.01, GraphPad Software Inc., La Jolla, CA, USA).

## Results

### Differences in average trait expression among populations

Expression of sociability and activity differed between populations, while boldness did not (‘population’ effect in Table 1; Fig. 3a). On a temporal scale, fish sampled at site 1 in August showed higher levels of sociability (i.e. reduced distance to robotic fish) compared to March and June populations (*post-hoc*, August site 1 *vs*. March – site 1: *p* = 0.003, *vs*. June – site 1: *p* = 0.025). Similarly, fish sampled at site 1 in August were more active than populations sampled in March and June (*post-hoc*, August site 1 *vs*. March – site 1: *p* = 0.002, *vs*. June – site 1: *p* < 0.001). On a spatial scale, we found no evidence that fish sampled in August at site 1 and site 2 differed in boldness, sociability, nor activity (*post-hoc*, August site 1 *vs*. site 2 – boldness: *p* = 0.76, sociability: *p* = 0.92, activity: *p* = 0.89; Fig. 3a).

**Table 1:** Results from linear mixed models.

| Effect | Boldness ($\log_{10}$) | | Sociability ($\log_{10}$) | | Activity | |
| --- | --- | --- | --- | --- | --- | --- |
| | $F_{df1,df2}$ | $p$ | $F_{df1,df2}$ | $p$ | $F_{df1,df2}$ | $p$ |
| population | $F_{3,469} = 0.57$ | 0.634 | $F_{3,469} = 3.8$ | **0.01** | $F_{3,469} = 8.44$ | **< 0.001** |
| trial | $F_{2,469} = 1.72$ | 0.18 | $F_{2,469} = 135.9$ | **< 0.001** | $F_{2,469} = 42.32$ | **< 0.001** |
| sex | $F_{1,469} = 0.06$ | 0.81 | $F_{1,469} = 2.28$ | 0.132 | $F_{1,469} = 0.24$ | 0.622 |
| body size (SL) | $F_{1,469} = 8.43$ | **0.004** | $F_{1,469} = 0.02$ | 0.884 | $F_{1,469} = 3.58$ | 0.059 |

For each behavioral trait, the LMM included 'population', 'trial' and 'sex' as fixed effects and 'body size' (SL in mm) as covariate with the random factor Fish ID. Significant effects ( $p < 0.05$ ) are indicated in bold.

Boldness and activity showed a size-dependency (sig. effect of covariate ‘body size’; Table 1, Fig. 3b) with smaller individuals being bolder (i.e. reduced emergence time; Pearson’s *r* = 0.32; *p* < 0.001) and more active (Pearson’s *r* = −0.35; *p* < 0.001) than larger ones. We found no such relation between body size and sociability (Pearson’s *r* = 0.002; *p* = 0.98). While sex is correlated with size in the sexually dimorphic guppy, sex did not affect any of the measured behavioral traits (non-sig. ‘sex’ effect; Table 1). We further observed a habituation to the repeated testing during sociability and activity assays (sig. ‘trial’ effect; Table 1). Fish were considerably less social and less active with each consecutive measurement.

### Among- and within-individual variation in behavioral traits

Fish within each sampled population differed consistently in their expression of boldness, sociability and activity across trials (sig. repeatability estimates; Table 2). Only fish sampled in August at site 2 had a significantly higher repeatability estimate for sociability than fish from the March sampling at site 1 (non-overlapping CIs; Table 2). There was no significant difference among populations in estimated overall behavioral variances in any of the examined traits (see ‘total variation’ in Table 2; for within- and among-individual variance estimates see Supplementary material Appendix 3, Table S4).

**Table 2:** Repeatability estimates and total behavioral variance estimates.

| Population | Boldness ($\log_{10}$) | | | Sociability ($\log_{10}$) | | | Activity | | |
| --- | --- | --- | --- | --- | --- | --- | --- | --- | --- |
| | Repeatability [95% CI] | $p$ | Total Variation [95% CI] | Repeatability [95% CI] | $p$ | Total Variation [95% CI] | Repeatability [95% CI] | $p$ | Total Variation [95% CI] |
| March site 1 | 0.49 [0.40–0.57] | **0.004** | 0.24 [0.15–0.4] | 0.26 [0.15–0.4] | 0.05 | 0.06 [0.04–0.1] | 0.27 [0.17–0.41] | **0.042** | 6.86 [4.3–11.87] |
| June site 1 | 0.40 [0.31–0.5] | **0.005** | 0.31 [0.2–0.5] | 0.32 [0.23–0.43] | **0.011** | 0.07 [0.05–0.11] | 0.30 [0.21–0.42] | **0.015** | 5.75 [3.74–9.33] |
| August site 1 | 0.54 [0.47–0.6] | **0.001** | 0.34 [0.22–0.54] | 0.34 [0.25–0.45] | **0.009** | 0.06 [0.04–0.09] | 0.38 [0.29–0.47] | **0.005** | 7.15 [4.61–11.48] |
| August site 2 | 0.53 [0.46–0.6] | **0.001** | 0.3 [0.19–0.48] | 0.49 [0.42–0.57] | **0.001** | 0.08 [0.05–0.13] | 0.36 [0.27–0.46] | **0.007** | 5.92 [3.81–9.57] |

### Behavioral syndrome structure

Boldness and activity were correlated in all four population samples (sig. correlation between boldness and activity: *F*_1,157_ = 52.23; *p* < 0.001; no sig. difference in slopes among populations: *F*_3,151_ = 0.45; *p* = 0.715). Fish with high average activity levels also left the shelter quicker, confirming that activity and boldness are part of a larger behavioral syndrome (Fig. 4).

**Figure 4:**
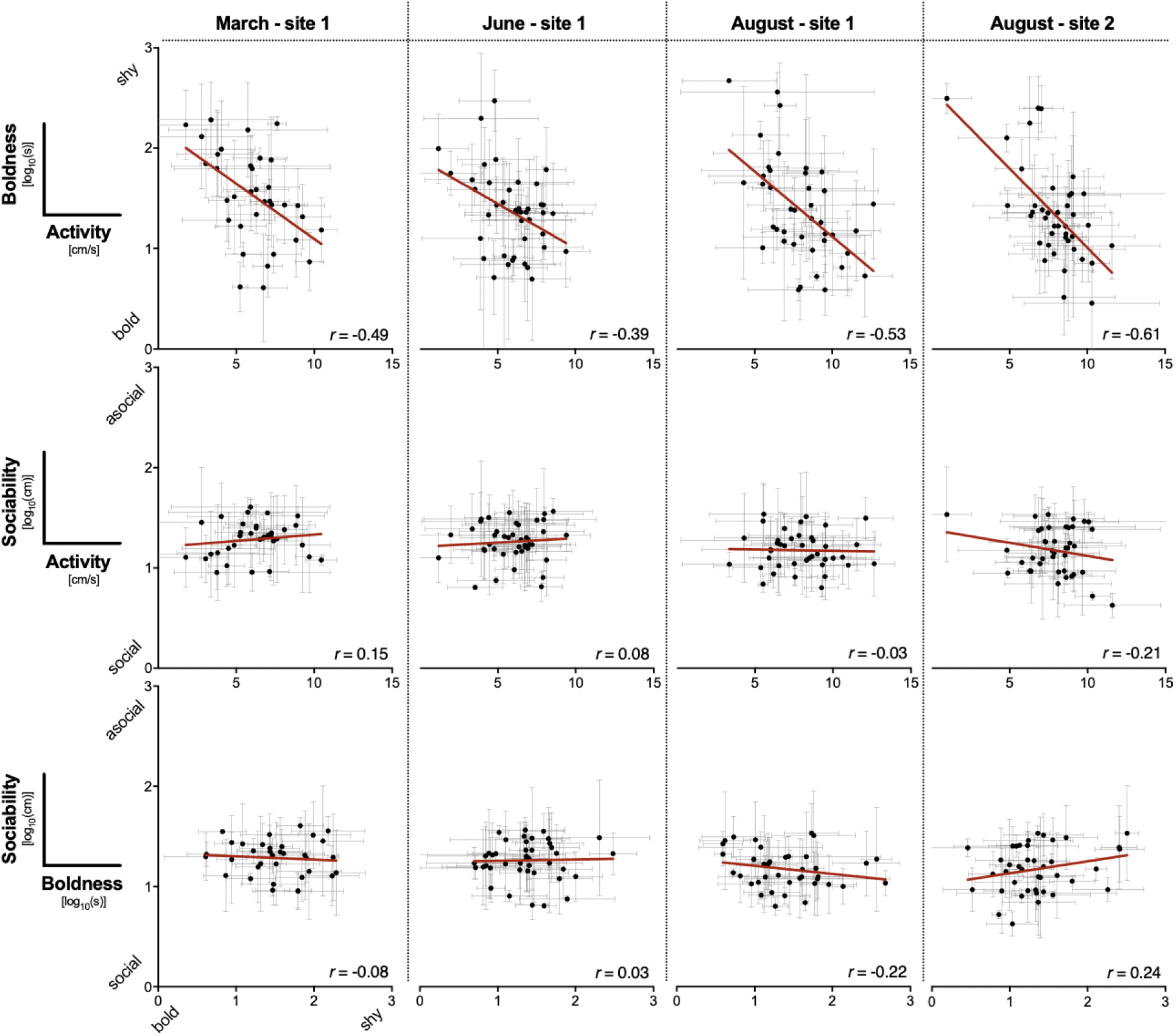
Correlations between behavioral traits across temporal and spatial scale. Mean trait scores* (±SD) of individual test fish are shown for each population. Linear regressions (red line and Pearson’s *r*) visualize the relationships between traits (see main text for detailed analysis of trait correlation; *n* = 159). * Note that for boldness and sociability lower values indicate a high trait score.

There was no evidence for correlations between sociability and boldness (*F*_1,157_ = 0.026; *p* = 0.872; non-sig. difference in slopes: *F*_3,151_ = 1.8; *p* = 0.149), nor between sociability and activity (*F*_1,157_ = 0.062; *p =* 0.803; non-sig. difference in slopes: *F*_3,151_ = 0.816; *p* = 0.487) among populations. Thus, sociability measured as distance kept to the moving robotic fish does not seem to be part of a behavioral syndrome with either boldness nor activity.

## Discussion

Our study follows the annual range shifts of non-native fish in an artificially heated stream in temperate Germany. As guppies dispersed into winter-abandoned areas in warmer months, we expected this process to be driven by dispersal-enhancing behavioral traits. We found similar consistent among-individual variation (e.g., significant behavioral repeatability) in the behaviors ‘boldness’, ‘sociability’ and ‘activity’ within all temporal and spatial samples (except a difference between sociability in March at site 1 and August at site 2). Contrary to our predictions, we did not detect a decrease in total behavioral variation of the sampled populations indicative of a “thinning out” of disperser’s behavioral types. Further, we found no evidence that the source and the newly-founded population differed in terms of average behavioral trait expression, though a seasonal change in activity and sociability (but not boldness) was detectable in the source population: August-sampled individuals were more active and more sociable than those sampled earlier in the year. Instead, we found a behavioral syndrome among activity and boldness that was consistently detectable throughout the year and across sampled populations. To sum up, we found no strong evidence for a personality-biased dispersal in this feral guppy population and the detected seasonal differences in average trait expression (sociability, activity) are likely due to changed environmental factors such as temperature, predator abundance and/or resource availability.

Differences in behavioral traits across seasons but not between sites as found in our study could be explained by seasonal variation in environmental conditions rather than by a drain of individuals carrying certain behavioral types from source to downstream populations. The Gillbach’s water temperatures show strongest differences between sites in the colder months but relatively small differences during the warmer months. Thus, when winter-abandoned peripheral sites are colonized again in summer, these habitats do not differ much in temperature from the source at this point in time. Nevertheless, the Gillbach still shows a strong seasonality in temperature which, in combination with a variation in day length, food quality/quantity and predation pressure, may likely cause population-wide changes in behaviors across seasons (e.g., Biro et al. 2010; Eccard and Herde 2013; Spiegel et al. 2015; Uchida et al. 2016; Barbosa et al. 2018; Dhellemmes et al. 2020) as seen in the source population. For example, prior temperature acclimation during sensitive developmental phases can alter the phenotypes of subsequent adult phases (Beitinger et al. 2000; Seebacher et al. 2014). This might explain the increased activity exhibited by fish in August, as higher developmental temperatures lead to more active phenotypes (Seebacher et al. 2014). However, teasing apart the effects of abiotic (e.g. temperature) and biotic (e.g., predation pressure) fluctuations in environmental conditions will require additional experimentation.

Besides seasonal differences in average behavioral traits, we found stable behavioral syndromes among sampled populations that involved a strong positive correlation between boldness and activity. Such a boldness-activity syndrome has been documented in various taxa (van Oers et al. 2004; Pintor et al. 2008; Wilson and Godin 2009; Cote, Fogarty, et al. 2010; Eccard and Herde 2013; Muraco et al. 2014), including guppies (Brown and Irving 2014). Natural selection through predation favors the evolution of behavioral correlations (Dingemanse et al. 2007; Harris et al. 2010; Dhellemmes et al. 2020). The Gillbach’s guppy population underwent strong seasonal changes in size and the observed population decline in summer, while primary production is highest in temperate regions, suggests strong predation pressure. A positive boldness-activity correlation might thus represent an optimal trait combination in the face of predation experienced by both populations (Smith and Blumstein 2010). However, the lack of differences in the level of (individual) behavioral variation among populations might be a result of a random downstream drift and a low overall behavioral plasticity in this population. This can be a consequence of the assumed bottleneck origin of this feral population that most likely stems from an unknown number of released domesticated guppies (see Gertzen et al. 2008 for estimates of propagule pressure for popular aquarium species). Investigations involving other populations of the guppy and more distantly related species are highly recommended, especially as several environmental factors are predicted to impact behavioral variation, especially in fish (e.g., Harris et al. 2010; Laskowski et al. 2016; Barbosa et al. 2018). Alternatively, guppies and especially those from the Gillbach that experience highly fluctuating temperature regimes may have the ability to adjust their behavioral expressions within very short times, e.g., within the 30-days acclimatization period in the laboratory used here. It is known that guppies have a remarkable capacity for behavioral plasticity (Deacon and Magurran 2016) and thermal adaptation (Chung 2001; Jourdan et al. 2014), and thus have successfully established populations in environments with very different (thermal) regimes compared to their native habitats (Deacon et al. 2011). Testing of individuals after different acclimation periods could be a possibility to investigate this issue further.

Theoretical models (Cote, Clobert, et al. 2010; Spiegel et al. 2017) as well as several studies across taxa (Duckworth and Badyaev 2007; Pintor et al. 2009; Cote, Fogarty, et al. 2010; Myles-Gonzalez et al. 2015; Coates et al. 2019) provide evidence for the prominent role of individual differences in behavior during dispersal and invasion processes (Bowler and Benton 2005; Chapple et al. 2012; Canestrelli et al. 2015). Nevertheless, the picture might be more complicated and context-dependent: spatial sorting of behavioral types did not emerge under predation pressure (Cote et al. 2013) and was lost as the population aged (Thorlacius et al. 2015). Moreover, even if there are significant (but small) effects of behavioural trait variation affecting relevant ecological phenomena, they might be non-detectable in benign laboratory environments (Schröder et al. 2016; Moiron et al. 2020). Our current study exemplifies that population-wide behavioral analysis can fail to provide evidence for behaviorally-mediated migration patterns and future analysis may focus on tracking individuals along their travel routes across seasons and locations. On top of this, several other feral guppy populations are known in Germany (Lukas, Kalinkat, et al. 2017) and across Europe (Kempkes et al. 2018) for which similar investigations are highly advisable in order to test the generality of our herein presented results. Such analyses should strive to incorporate the invasion history, environmental factors as well as the population-specific genetic backgrounds which all can affect behavioral variation in wild populations (Sih and Bell 2008; Bergmüller and Taborsky 2010; Stamps and Groothuis 2010).

## Notes

### Competing Interest Statement

The authors have declared no competing interest.

