## Supplemental for "Consistent behavioral syndrome across seasons in an invasive freshwater fish"

### Supplement

#### 1. Fish community

Samplings revealed 13 different fish taxa (capture or visual record, see Table S1), seven of which were non-native or ornamental species. Guppies were collected for behavioral testing, while ‘bycatch’ (i.e. all other native or non-native fish captured) were immediately euthanized with an overdose of clove oil and preserved in 99% ethanol for subsequent identification. Specimens were determined to the lowest feasible taxonomic level using the most recent literature available (Trewavas 1983; Teugels & Thys Van den Audenaerde 1992, 2003; Kottelat & Freyhof 2007; Schmitter-Soto 2007a,b; Stiassny *et al*. 2007) and have been integrated into the collection of the Zoological Research Museum Alexander Koenig.

Among the non-native species, the guppy *Poecilia reticulata* was the most abundant. Guppies were present at the Gillbach’s source (site 1) and one kilometer downstream (site 2) during all three samplings, however, we did not encounter any guppies beyond site 2. Notably, we found newly born fry and juveniles during all samplings, indicating that reproduction occurs year-round. Further, the feral guppy population of the Gillbach showed a high plasticity in body coloration and fin dimension ranging from the sexual dimorphism typically encountered in natural guppy populations to the ornamentation derived from various strains used in the aquarium trade (Fig. S2).

**Table S1: Fish community at the thermally-altered Gillbach** Summary of native and non-native fish observed (V) or caught (X) at the source of the Gillbach (SI), one kilometer downstream (SII) or further down (FD) during three samplings in 2016. Note that sampling efforts were not standardized for examining species composition (i.e. sampling ceased once a sufficient number of adult *Poecilia reticulata* for experimental purposes were caught; see Table S3) and thus includes records of species previously described for this system.

|  | Species | Origin | March | | | June | | | August | | | Previous records |
| --- | --- | --- | --- | --- | --- | --- | --- | --- | --- | --- | --- | --- |
|  |  |  | SI | SII | FD | SI | SII | FD | SI | SII | FD |  |
| Anguillidae | *Anguilla anguilla* (Linnaeus, 1758) | Europe |  |  |  |  |  |  |  |  |  | 3 |
| Cichlidae | *Amatitlania nigrofasciata* (Günther, 1867) | Central America | X | X |  | X | X | X | X | X | X | 2,3,4,5,8 |
|  | *Maylandia aurora* (Burgess, 1976) | Africa |  |  |  |  |  |  |  |  |  | 3 |
|  | *Oreochromis* sp. | Africa |  | X | V |  |  | X | X |  | V | 3,4,5,8 |
|  | *Pelmatolapia mariae* (Boulenger, 1899) | Africa |  |  |  |  |  |  |  |  | V | 8 |
| Cyprinidae | *Barbus barbus*  (Linnaeus, 1758) | Europe |  |  |  | X | X |  |  |  |  | 3,4,8 |
|  | *Carassius auratus (gibelio)* (Linnaeus, 1758) | ornamental | V |  |  | V |  |  |  |  |  | 3,4 |
|  | *Chondrostoma nasus* (Linnaeus, 1758) | Europe |  |  |  |  |  |  |  |  | V | 4 |
|  | *Cyprinus carpio*  (Linnaeus, 1758) | Europe/Asia |  |  |  |  |  | V |  |  | V | 4 |
|  | *Gobio gobio*  (Linnaeus, 1758) | Europe |  | X |  |  |  |  |  | X | X | 2,3,4 |
|  | *Pseudorasbora parva* (Temminck & Schlegel 1846) | Asia |  |  |  |  |  |  |  |  |  | 3,4 |
|  | *Squalius cephalus*  (Linnaeus, 1758) | Europe | X | X |  | X | X | X | X | X | X | 2,3,4,8 |
| Loricariidae | *Ancistrus* sp. | South America |  |  |  |  | X | V | X |  | X | 4,5 |
| Poeciliidae | *Poecilia reticulata* (Peters, 1859) | South America | X | X |  | X | X |  | X | X |  | 3,4,5,7,8 |
|  | *Poecilia sphenops*  (Valenciennes, 1846) | Central/South America | V |  |  |  |  |  |  |  |  |  |
|  | Poeciliidae I |  |  |  |  |  |  |  |  |  |  | 3 |
|  | Poeciliidae II |  |  |  |  |  |  |  |  |  |  | 3 |
| Siluridae | *Silurus glanis* (Linnaeus, 1758) | Europe |  |  |  |  |  |  |  |  | V |  |

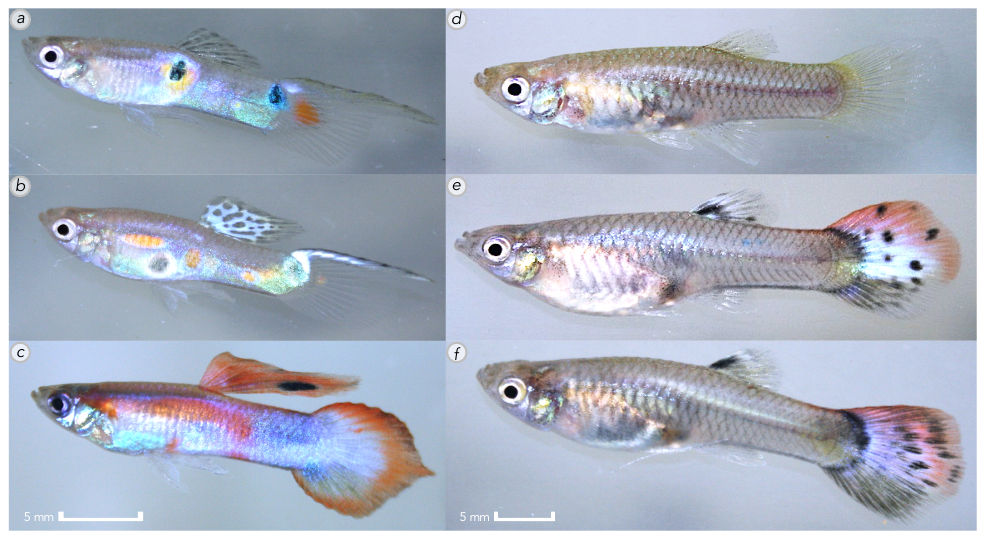

**Figure S2: Specimens of feral guppies (*Poecilia reticulata*) collected from the Gillbach population in 2016.** Wild-type guppies are sexually dimorphic with males displaying vivid orange, black and green spots or stripes (a), while females are colored an inconspicuous grey or brown (d). In the Gillbach population, both males (a–c) and females (d–f) show ­polymorphism in coloration and fin structure ­– including some rare color morphs (c, e, f) – typical for fish of the ornamental trade.

#### 2. Size estimation of the source population (site 1)

To estimate changes in population size as well as the degree of philopatry of the source population, we marked and released guppies sampled at site 1 during June and August of 2016. Visible implant elastomer tags (Northwest Marine Technology Ltd) were injected subcutaneously (for procedure details see Jourdan et al. 2014) using two colors (i.e. pink and green, respectively) to clearly differentiate between capture events. No previously marked fish were marked a second time. Recapture rates were recorded during subsequent samplings and analyzed with a log-linear model for closed populations (Mt in package *Rcapture*; Baillargeon and Rivest 2007) after Jourdan et al. (2014).

Overall, we tagged 150 guppies during two samplings and recovered 23 marked individuals (9 June tags retrieved 86 days post-release and 14 August tags retrieved 22 days post-release; Table S3). We estimated a total population size of 4487 (95% CI: 2584.2–9108.7) adult guppies in June and 514 (95% CI: 351.7–846.1) in August. The number of guppies that were taken to the laboratory for subsequent behavioral testing was added to the estimates, resulting in a population size estimate of about 4584 individuals in June and 688 individuals in August.

**Table S3: Sampling results of guppies at the Gillbach including total captures and number of marked and recaptured individuals.**

| **Sample** | | **# Captured** | | | | **# Tagged** | **# Recaptured** |
| --- | --- | --- | --- | --- | --- | --- | --- |
|  |  | **Total** | **Male** | **Female** | **Juvenile** |  |  |
| **March** | SI | 54 | 10 | 25 | 19 | - | - |
|  | SII | 11 | 3 | 5 | 3 | - | - |
| **June** | SI | 243 | 83 | 113 | 47 | 146 | - |
|  | SII | 4 | 1 | 1 | 2 | - | - |
| **August** | SI | 278 | 98 | 121 | 59 | 104 | 9 |
|  | SII | 65 | 19 | 33 | 13 | - | 0 |
| **September** | SI | 70 | 17 | 50 | 3 | - | 14 |
|  | SII | - | - | - | - | - | - |

#### 3. Behavioral testing

**Table S4: Variances from Repeatability models.** Shown are estimates of within- and among-individual variance along with 95% confidence intervals computed from linear mixed models (see main text) for each behavioral traits of all population samples.

|  | Boldness (log_10_) | | Sociability (log_10_) | | Activity | |
| --- | --- | --- | --- | --- | --- | --- |
| Population | *Var*_within_  [95% CI] | *Var*_among_ [95% CI] | *Var*_within_  [95% CI] | *Var*_among_ [95% CI] | *Var*_within_  [95% CI] | *Var*_among_ [95% CI] |
| March – site 1 | 0.12  [0.09–0.17] | 0.12  [0.06–0.23] | 0.044  [0.031–0.061] | 0.015  [0.006–0.041] | 5.00  [3.58–6.98] | 1.87  [0.71–4.89] |
| June – site 1 | 0.19  [0.14–0.25] | 0.12  [0.06–0.25] | 0.047 [0.035–0.064] | 0.022  [0.010–0.048] | 4.03  [2.97–5.46] | 1.73  [0.77–3.88] |
| August – site 1 | 0.16  [0.12–0.21] | 0.18  [0.1–0.32] | 0.037  [0.028–0.051] | 0.019  [0.009–0.041] | 4.46  [3.28–6.06] | 2.69  [1.33–5.43] |
| August – site 2 | 0.14  [0.1–0.19] | 0.16  [0.09–0.29] | 0.043  [0.031–0.058] | 0.041  [0.023–0.075] | 3.79  [2.78–5.16] | 2.14  [1.04–4.41] |
